# Comparing surrogates to evaluate precisely timed higher-order spike correlations

**DOI:** 10.1101/2021.08.24.457480

**Authors:** Alessandra Stella, Peter Bouss, Günther Palm, Sonja Grün

**Affiliations:** Institute of Neuroscience and Medicine (INM-6) and Institute for Advanced Simulation (IAS-6) and JARA Institute Brain Structure-Function Relationships (INM-10), Jülich Research Centre, Jülich, Germany; Institute of Neural Information Processing, University of Ulm, Ulm, Germany; Theoretical Systems Neurobiology, RWTH Aachen University, Aachen, Germany

## Abstract

The generation of surrogate data, i.e., the modification of original data to destroy a certain feature, is used for the implementation of a null-hypothesis whenever an analytical approach is not feasible. Thus, surrogate data generation has been extensively used to assess the significance of spike correlations in parallel spike trains. In this context, one of the main challenges is to properly construct the desired null-hypothesis distribution and to avoid a bias in the null-hypothesis by altering the spike train statistics.

A classical surrogate technique is uniform dithering (UD), which displaces spikes locally and uniformly. In this study, we compare UD against five surrogate techniques (two newly introduced) in the context of the detection of significant spatio-temporal spike patterns. We evaluate the surrogates for their performance, first on spike trains based on point process models with constant firing rate, and second on modeled non-stationary artificial data serving as ground truth to assess the pattern detection in a more complex and realistic setting. We determine which statistical features of the original spike trains are modified and to which extent. Moreover, we find that UD fails as an appropriate surrogate because it leads to a loss of spikes in the context of binning and clipping, and thus to a large number of false-positive patterns. The other surrogates achieve a better performance in detecting precisely timed higher-order correlations. Based on these insights, we analyze experimental data from pre-/motor cortex of macaque monkeys during a reaching-and-grasping task for spatio-temporal spike patterns.

**Significance statement:** Temporal jittering or dithering of single spikes or subsections of spike trains is a common method of generating surrogate data for the statistical analysis of temporal spike correlations. We discovered a serious problem with the classical and widely used method of uniform dithering that can lead to an overestimation of significance, i.e., to false positives in the statistical evaluation of spatio-temporal spike patterns. Therefore we consider 5 other dithering methods, compare and evaluate their statistical properties. Finally, we apply a much better method (trial shifting) to the analysis of experimental multiple-unit recordings and find several highly significant patterns that also reflect different experimental situations.

## 1 Introduction

The usage of surrogates has become a standard tool in data analysis and computational statistics (Louis et al., 2010b; Abeles and Gat, 2001; Grün et al., 2002a; Grün, 2009). It is often used to replace the definition of an appropriate null-hypothesis in classical statistical testing, which describes the possibility that the observed effects or results have occurred merely by chance. In statistical textbooks and software packages, standard definitions of null-hypotheses are used, which allow an analytical derivation and automatic computation of the corresponding significance probabilities. In computational statistics, it has become possible to determine these probabilities also for more complex and not analytically tractable null-hypotheses, by extensive sampling from the null-hypothesis distribution, i.e., by generating artificial data from this distribution. The generation of surrogate data follows a completely different approach. In exploratory data analysis or scientific investigations, we may have observed an interesting effect, but we often have no idea what would be an appropriate model for “randomness”, and it would be premature to assume a standard random model, like the normal distribution, just for convenience. In this situation, we can use the data themselves to test for the significance of the observed effect by simply modifying them to generate more data of the same type, which can then be used to determine significance probabilities. Such methods are called bootstrapping (Efron and Tibshirani, 1993) and typical methods are resampling from or reordering of the data, or adding small amplitude noise. The generation of surrogate data is a particular version of this, which is typically used when we have an idea or hypothesis concerning the features in the data that are relevant for the effect. In this case, we modify or add some noise to the data in order to destroy these features. If we use these modified data as our “null-hypothesis” and the observed effect does not occur or occurs with a very low probability, we have obtained evidence that those features are indeed relevant and our hypothesis was correct (Kass et al., 2005; Ventura, 2010).

Here we are interested in the interactions between hundred or more neurons that were recorded in parallel by multiple electrodes (Riehle et al., 2013; Brochier et al., 2018). In view of the apparent randomness in the reaction of neurons to repeated stimuli (Nawrot et al., 2008) one important question concerns the temporal precision of neural interactions, which has been studied by means of pairwise (Grün et al., 2002a,b; Pipa and Grün, 2003; Pipa et al., 2003, 2007; Grün, 2009; Pipa et al., 2013) and higher-order correlation analysis (Villa and Abeles, 1990; Martignon et al., 1995; Riehle et al., 1997; Prut et al., 1998; Villa et al., 1999; Kilavik et al., 2009; Shimazaki et al., 2012) of parallel recorded multiple-unit spike trains. In order to demonstrate high precision in temporal multiple-neuron interactions by statistical methods, one needs surrogate methods that destroy correlations at a high temporal precision, but not at a low temporal precision and maintain as much as possible all other statistical properties of the individual spike trains. So, the basic idea was to shift the spikes of the original data by a small amount. These surrogate methods are called dithering or jittering or teetering (Date et al., 1998; Hatsopoulos et al., 2003; Pazienti et al., 2008; Stark and Abeles, 2009; Louis et al., 2010b); the most commonly used of these methods is uniform dithering, which shifts each individual spike by a small uniformly distributed amount. Unfortunately, this method suffers from severe problems concerning the preservation of statistical properties like the inter-spike interval distribution and can lead to substantial spike count reduction when combined with binning and binarization of spike trains, which is a prerequisite for many methods of statistical analysis. Consequently, we introduced some different surrogate methods and compared them with each other and with uniform dithering in terms of statistical properties, and finally also the effect on significance evaluation of repeating spatio-temporal spike patterns (SPADE; Torre et al., 2013; Quaglio et al., 2017; Stella et al., 2019). The comparison is done on artificially generated data (both stationary and non-stationary) and also on experimental multiple-unit recordings, where we find some interesting highly significant spike patterns. It turns out that only uniform dithering can lead to an overestimation of pattern significance, i.e. to false positive detection of patterns.

## 2 Materials and Methods

### 2.1 Surrogate methods

Different types of surrogates were already developed, however, in the context of different analysis methods (Louis et al., 2010a,b; Grün, 2009; Pipa et al., 2008, 2013). Here, we compare six different surrogate methods, four known from the literature and two newly developed by us, and evaluate their applicability for significance assessment in spike train correlation analyses. Since we give particular attention to surrogate techniques that are supposed to preserve as many features of the original spike train as possible, we do not consider surrogates that destroy the firing rate profile.

#### 2.1.1 Uniform dithering

The uniform dithering method consists in displacing each individual spike of each neuron by a small uniformly distributed random jitter ~ *U* [-Δ, +Δ] around its original position. An example sketch is shown in Fig 1A. It is also known by the names: jittering or teetering and is a classical choice for surrogate generation and was employed in several experimental studies (Abeles and Gat, 2001; Hatsopoulos et al., 2003; Gerstein, 2004; Shmiel et al., 2006; Maldonado et al., 2008; Torre et al., 2016). Due to its simplicity and computational speed, it is widely used for detection of pairwise synchrony (i.e., cross-correlogram significance estimation; Grün, 2009; Louis et al., 2010b), higher-order synchrony, and pattern detection (Abeles and Gat, 2001; Gansel and Singer, 2012; Torre et al., 2016). In particular, it was chosen as the surrogate generation technique for synchrony and pattern detection using SPADE (Torre et al., 2016; Quaglio et al., 2017; Stella et al., 2019). However, Louis et al. (2010a) already demonstrated in the context of pairwise synchrony analysis that uniform dithering (UD) can lead to false positives for regular firing properties (CV *<* 1).

**Figure 1.**
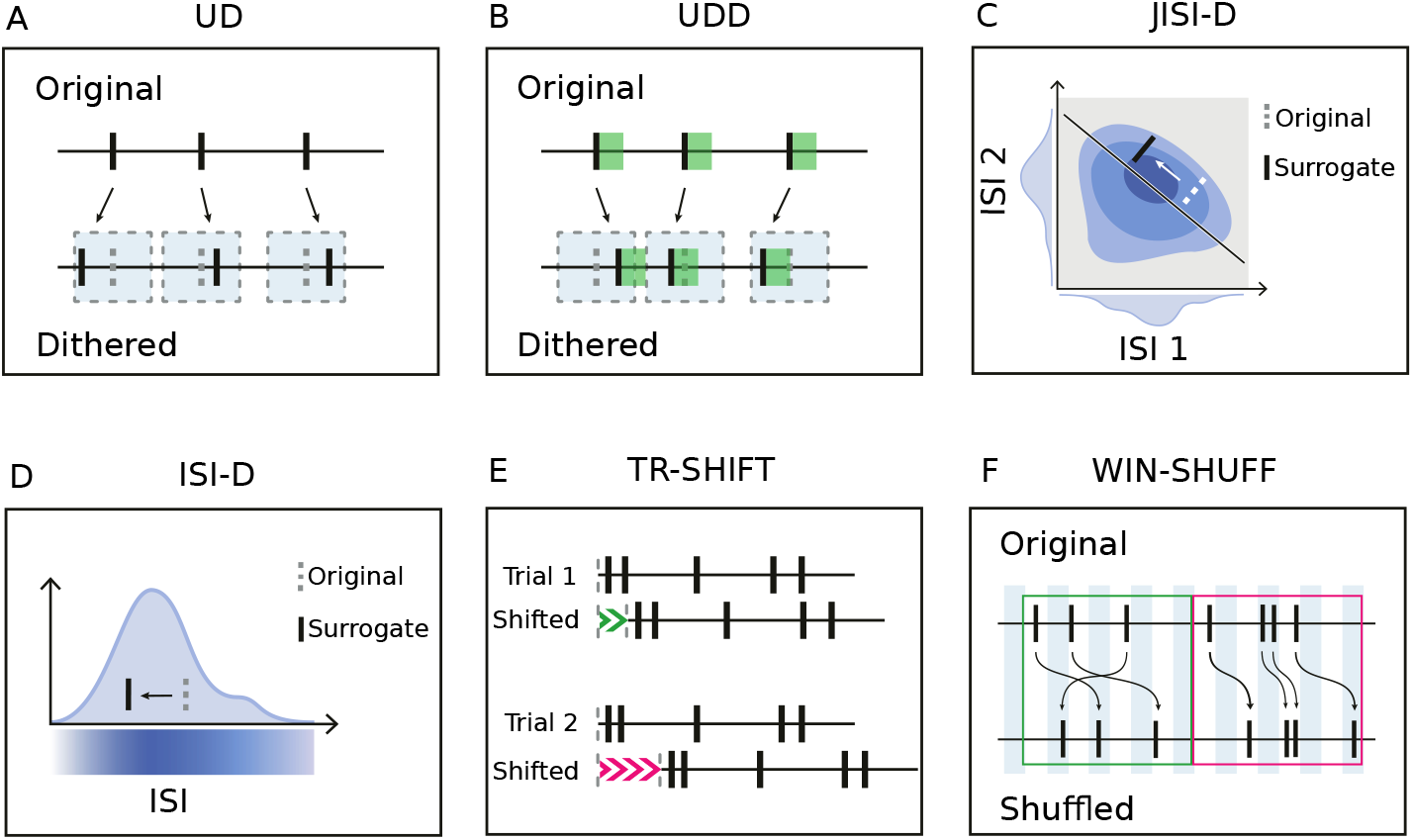
Sketches of the surrogate techniques. (A). Uniform Dithering (UD). Each spike is displaced according to a uniform distribution centered on the spike. Grey dotted rectangles represent the dithering window. (B). Uniform Dithering with dead-time (UDD). Similar to uniform dithering, but spikes are constrained not to be closer to each other than a given dead-time. Green shadows represent the dead-time after each spike. Grey dotted rectangles represent the dithering window. (C). Joint Inter-Spike Interval Dithering (JISI-D). Each spike is displaced according to the J-ISI distribution of the neuron, sampled from the data. J-ISI distribution, projected in a two-dimensional plane, is represented in blue. On the *x* and *y*-axes we represent the projections of the first and second ISI given three spikes. (D). Inter-Spike Interval Dithering (ISI-D). Each spike is displaced according to the ISI distribution of the neuron, sampled from the data. The ISI distribution is represented in blue, along with its intensity. (E). Trial Shifting (TR-SHIFT). Each trial is shifted by a randomly chosen amount from a uniform distribution (represented in green and pink), independently across trials and neurons. (F). Window Shuffling (WIN-SHUFF). Binned spike data are shuffled within exclusive windows (marked green and pink).

The dither parameter of the method Δ *>* 0 determines the maximal displacement of a spike from its original position. It needs to be selected appropriately, e.g., in the range of 15ms to 25ms (Stella et al., 2019; Torre et al., 2016), and is typically a multiple of the bin size parameter. If Δ is too small, it causes insufficient displacement of the correlated spikes and may lead to an underestimation of significance, whereas if Δ is too large, it yields a strongly smoothed firing rate profile (Pazienti et al., 2008) and, thus, an inappropriate null-hypothesis.

#### 2.1.2 Uniform dithering with dead-time

We introduce uniform dithering with dead-time (UDD) as a variant of the classical uniform dithering (Fig 1B). As the name suggests, the estimated dead-time *d* is conserved during the temporal displacement of each spike. This is done by limiting the window into which spikes can be dithered. The uniform displacement of each spike is limited by the intervals to the neighboring spikes minus the dead-time *d*. Thus, it does not allow two dithered spikes to have a temporal distance smaller than the dead-time, and, unlike for UD, the displacement of each spike is hence not independent of its neighbor. As described later in section 3.2.1, a dead-time may be introduced by spike sorting. Further, the biological absolute refractory period of neurons can yield minimal intervals larger than those inserted by the spike sorting. We estimate the dead-time for each neuron to be the minimum inter-spike interval across all trials. In case of low firing rates, the minimum ISI may be still in the range of hundreds of milliseconds, complicating the estimation of a biologically reasonable dead-time. For this reason, we define a maximal dead-time parameter *d_max_* such that, if the minimal inter-spike interval exceeds *d_max_* (in our case 4ms), then we set *d* = *d_max_*.

#### 2.1.3 Joint-ISI dithering

Gerstein (2004) suggested to dither spikes according to their joint-ISI distribution of adjacent ISI intervals. Joint inter-spike interval dithering (JISI-D; Fig 1C; Gerstein, 2004; Louis et al., 2010a) aims to keep the distribution of the preceding and following inter-spike intervals relative to a spike according to the joint-ISI probability distribution. This probability distribution is derived for each neuron from its spike train by calculating the corresponding joint-ISI histogram (with a default bin size of 1ms). Dithering one spike according to such a two-dimensional histogram corresponds to moving the spike along the anti-diagonal of the joint-ISI distribution (Gerstein, 2004; Louis et al., 2010a).

Unfortunately, the recordings are often non-stationary and too short to comprise enough spikes to estimate the underlying joint-ISI probability distribution. Therefore, we apply on the joint-ISI histogram a 2d-Gaussian smoothing with variance *σ*^2^, with *σ* of the order of milliseconds (Louis et al., 2010a).

#### 2.1.4 ISI dithering

ISI dithering (ISI-D; Fig 1D), unlike JISI-D, does not consider the pair of a current and its subsequent ISI, instead, it dithers the individual spikes according to the ISI probability distribution. However, for practical reasons, we implemented ISI-D as a special case of the joint-ISI dithering assuming that two consecutive ISIs are independent, i.e., that the joint-ISI histogram can be represented as the product of the ISI histogram with itself, (*p*j-ISI(*τ, τ′*) = *p*ISI(*τ*) *p*ISI(*τ′*)). Thus, in comparison to the joint-ISI dithering, ISI-D does not take into account the correlations of subsequent ISI pairs and is particularly useful when there are not enough data to estimate the joint-ISI distribution. As a result, not the distribution of pairs of subsequent ISIs are preserved, but the distribution of the single ISIs irrespective of their order.

#### 2.1.5 Trial shifting

An alternative to the dithering of single spikes is dithering the entire spike train. Trial shifting (TR-SHIFT; Fig 1E; Pipa et al., 2008; Louis et al., 2010b) consists of shifting all spike times identically by a random uniform amount ~*U* [−Δ, +Δ], independently neuron by neuron and trial by trial. The method requires the time randomization to be different across neurons in the different trials, such that repeating identical patterns are shifted into different patterns from trial to trial. Therefore the method requires a segmentation of the spike trains into longer temporal sequences, called trials here. These could also be longer spike sequences that are separated by relatively long intervals between spikes as suggested by Harrison and Geman (2009). In our case, the trials are defined by the experimental protocol. TR-SHIFT has the benefit of keeping the entire spike train structure intact during each segment.

#### 2.1.6 Window shuffling

We also introduce window shuffling (WIN-SHUFF; Fig 1F), which divides the spike train into successive and exclusive small windows of predefined duration ΔWS, and further divides the windows into bins of length *b* (ΔWS should then be a multiple of *b*). The bins are then shuffled within each window. Additionally, spike times are randomized within each bin. The firing rate profile is modified by the local shuffling of the spikes to be stationary inside in window of duration ΔWS. To facilitate the comparison to the other methods, we fix throughout the paper ΔWS = 2Δ.

### 2.2 SPADE

We aim to apply the surrogates to the spike correlation method called Spike PAttern Detection and Evaluation (SPADE; Stella et al., 2019), which is capable of detecting spatio-temporal spike patterns across neurons and evaluates their significance.

The spike train data are first discretized into exclusive time intervals (bins). Typically, the bin length consists of a few milliseconds, which at the same time defines the allowed temporal imprecision of the neuronal coordination. The procedure of discretization counts the number of spikes within each bin (binning, Grün et al., 2002a; Torre et al., 2013), followed by reducing the bin content to 1 if a bin contains more than 1 spike (clipping). In the following, we will call the combination of these two steps *binarization*. Candidate spatio-temporal patterns are then mined using the Frequent Itemset Mining (FIM) algorithm (Zaki, 2004; Borgelt, 2012; Picado-Muiño et al., 2013), which yields the number of occurrences of each non-trivial detected spike pattern, along with its occurrence times. A non-trivial pattern is defined as one that, repeats at least a fixed number of times but cannot be explained as part of a larger pattern. The binarized data are the input to the mining algorithm. Pattern counts are then collected in the so-called *pattern spectrum*, i.e., the pattern counts are entered in a 3d-histogram according to their number of spikes *z*, the number of pattern repetitions *c*, and the temporal extent from first to last spike *d* (see Fig 5C). The triplet {*z, c, d*} is called *signature*.

FIM efficiently collects and counts pattern candidates, nonetheless, the statistical significance of each of the mined patterns has still to be evaluated. SPADE aims at testing whether the patterns emerge as an effect of precisely timed neuronal coordination, or merely by the firing rate of independently firing neurons. For this reason, the null-hypothesis is that spike trains are mutually independent given their firing rate (co-)modulations and that the occurring patterns in the data are given by chance. All patterns that have a p-value lower than the threshold (after multiple testing correction) reject the null-hypothesis and are classified as significant.

SPADE uses surrogate generation to implement the null-hypothesis, and, the standard method is UD. To generate reliable statistics, the method generates and analyzes many surrogates (in the range of thousands: here 5000), and each surrogate data set is subject to the same procedure as the original data: binarization, followed by FIM, which results in pattern counts per signature. From the pattern counts the p-value spectrum is calculated, i.e., a 3-dimensional matrix containing for each signature the fraction of surrogates data sets containing patterns with that signature, thus, a p-value for each signature (Fig 5C). The significance level is corrected by the False Discovery Rate correction (Benjamini and Hochberg, 1995), where the number of tests is the number of occupied entries of the pattern spectrum having the highest number of occurrences per size and duration. The significance evaluation step is called *pattern spectrum filtering* (PSF) which retrieves potentially significant patterns, that are further filtered by the *pattern set reduction* (PSR; Torre et al., 2013). Pattern set reduction consists of conditional tests of each pattern given any other pattern surviving the PSF, in order to remove spurious false positives resulting as a by-product of the overlap of true patterns and chance spikes.

When analyzing large-size experimental data, the search for all possible patterns can result in obtaining millions, if not billions, of putative patterns. Thus, the analysis can quickly become infeasible due to memory or time requirements. To optimize the analyses, we run the FIM algorithm separately per pattern size (from size 2 to size 10+ in steps of one). For each pattern size, we set a minimum number of occurrences *min_occ_* a pattern has to occur to be further considered after the mining step. This parameter is a rough estimate of the number of patterns expected by chance assuming independence of the spike trains, using stationary Poisson processes with the average rate of each of the spike trains for this estimation. Patterns with a lower number of occurrences are considered as spurious as they would be rejected anyway by the following statistical test. In addition, we fix the minimum number of occurrences of a pattern to be at least 10, i.e., 30% of the number of trials of the considered experimental data. The FIM output is then aggregated for the pattern spectrum filtering and the pattern set reduction.

Finally, as a result of the explained steps, SPADE outputs significant STPs, together with their number of occurrences, the involved neurons, the lags between the pattern spikes, and the times of pattern occurrences.

### 2.3 Experimental data and preprocessing

In the results section of the paper, we make use of experimental data in two respects: a) we simulate independent data to test for false positives, and therefore extract statistical features of the experimental data to be included in the data; b) we analyze the experimental data for spike correlation by the usage of the various surrogates. The experimental data were recorded during a reaching-and-grasping task from the pre-/motor cortex of two macaque monkeys. Both monkeys were chronically implanted with a 100-electrode Utah array (Blackrock Microsystems, Salt Lake City). The experimental protocol is schematized in Fig 3A and was also published in Brochier et al. (2018) and in Riehle et al. (2013). Monkeys N and L were trained to self-initiate a trial by pressing a start button (registered as trial start, TS). Then, after a fixed time of 400ms, a visual signal (yellow LED) was shown, to attract the attention of the monkey (waiting signal, WS). After another 400ms-long waiting period, a first visual cue (2 LEDs on) was presented to the monkey for a period of 300ms (from CUE-ON to CUE-OFF) indicating the grip type: full-hand side grip (SG) or two-fingers precision grip (PG). Followed by another waiting period of 1000ms, the GO-signal was presented, containing also the information of the expected grip force (high, HF, or low, LF, by LEDs). The behavioral conditions were selected in a randomized fashion. The start of the movement of the monkey was recorded as the release of a switch (SR). Subsequently, the object touch (OT) and the beginning of the holding period (HS) are indicated. After 500ms of holding the object in place, a reward (RW) was given to the monkey and the trial finished.

The experimental data sets consist of two sessions (i140703-001 and l101210-001) of 15min of electrophysiological recordings containing around 35 trials per trial type, i.e., combinations of grip and force type. Each session is spike sorted using the Plexon Offline Spike Sorter (version 3.3). The spike data were extracted and are available on https://gin.g-node.org/INT/multielectrode_grasp.

We only consider neurons satisfying the following constraints: SNR *<* 2.5 (signal-to-noise ratio of spike shapes), average firing rate across trials *<* 70Hz. Hyper-synchronous (artifact) spikes across electrodes arising at the sampling resolution are detected automatically, classified as artifacts, and finally removed as in Torre et al. (2016). Only successful trials are retained. The experimental sessions are analyzed separately. For the analysis, trials are segmented into six 500ms-long separate epochs, to account for the behaviorally relevant events (as in Torre et al., 2016, and represented in Fig 3B). Segments of the same epochs in the same trial type are concatenated and yield 24 (4 trial types ×6 epochs) data sets per session.

### 2.4 Artificial non-stationary data

The simulated artificial data sets consist of non-stationary spike trains, with similar single spike-train features as the experimental data but without precise time correlations. This is done by generating as many spike trains as in the concatenated experimental data using the original firing rate profiles of the individual neurons, estimated with an optimized kernel density estimation as designed in Shinomoto (2010) and Shimazaki and Shinomoto (2010) on a single trial-by-trial basis. To account for dead-time and regularity of the data, individual spike trains are modeled separately as a Poisson process with dead-time (PPD; Deger et al., 2012) and as a Gamma process. Mathematically these processes are defined only in the stationary case. Since the time-varying firing rates are an important property of the experimental data which has to be incorporated in any “null-hypothesis”, we had to find a way of generating non-stationary spike data with the required ISI regularities. The PPD is a variation of the classical Poisson process wherein no spike is generated within an interval to the previous spike smaller than the dead-time *d*. The Gamma process has a shape parameter *γ* which is related to the intrinsic regularity/ irregularity of the spike train (Nawrot et al., 2008). If *γ* > 1, the process is regular (i.e., CV *<* 1); if instead *γ* = 1, it coincides with the classical Poisson process with an exponential ISI distribution. We do not consider Gamma processes with *γ* < 1 (CV *>* 1), i.e., bursty spike trains because our data did not contain such cases.

For the PPD data, we estimate the dead-time for each neuron and each combination of epoch and trial type by taking their minimum ISI. For the data modeled as a Gamma process, we instead fix the shape factor for each neuron and each combination of epoch and trial type by estimating the CV of the process in operational time (Nawrot et al., 2008) and then transform the CV into the shape factor (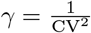, van Vreeswijk, 2010). The process is generated in operational time and then transformed back into real time. We then evaluate regularity through the CV2 measure, which compensates for non-stationary firing rates (Holt et al., 1996). The resulting CV2 distribution of all neurons of the data, simulated as a Gamma process, is very close to the one of the experimental data (Fig 8A, third inset). Note that the Gamma process does not have an absolute dead-time but for *γ* < 1, it has a low probability for small ISIs and can be regarded as containing a relative dead-time (Nawrot et al., 2008). The resulting firing rates of the artificial data are, for both generative models, close to the ones of the original spike trains.

### 2.5 Code accessibility

The code to perform and reproduce the analyses presented in this study can be found at (https://github.com/INM-6/SPADE_surrogates), along with the code to reproduce the figures contained in this paper, i.e., Fig 3B, Fig 4, Fig 5, Fig 6, Fig 7, Fig 8, Fig 9, Fig 10. Fig 1, Fig 2, Fig 3A are sketches created manually with vector graphics editors. The experimental data analyzed in section 3.4 can be found at https://gin.g-node.org/INT/multielectrode_grasp. The artificial data are generated from the experimental data within the *SPADE_surrogates* repository. The SPADE method and all implementations of the surrogate techniques are included in the Elephant Python package http://python-elephant.org. Regarding the computational cost, several improvements have been made for the performance of both, SPADE and the surrogate implementations (https://elephant.readthedocs.io/en/latest/release_notes.html; Porrmann et al., 2021). Nonetheless, depending on the size of the data set and the number of surrogates employed, large analyses can still take up to several hours on a computer cluster.

**Figure 2.**
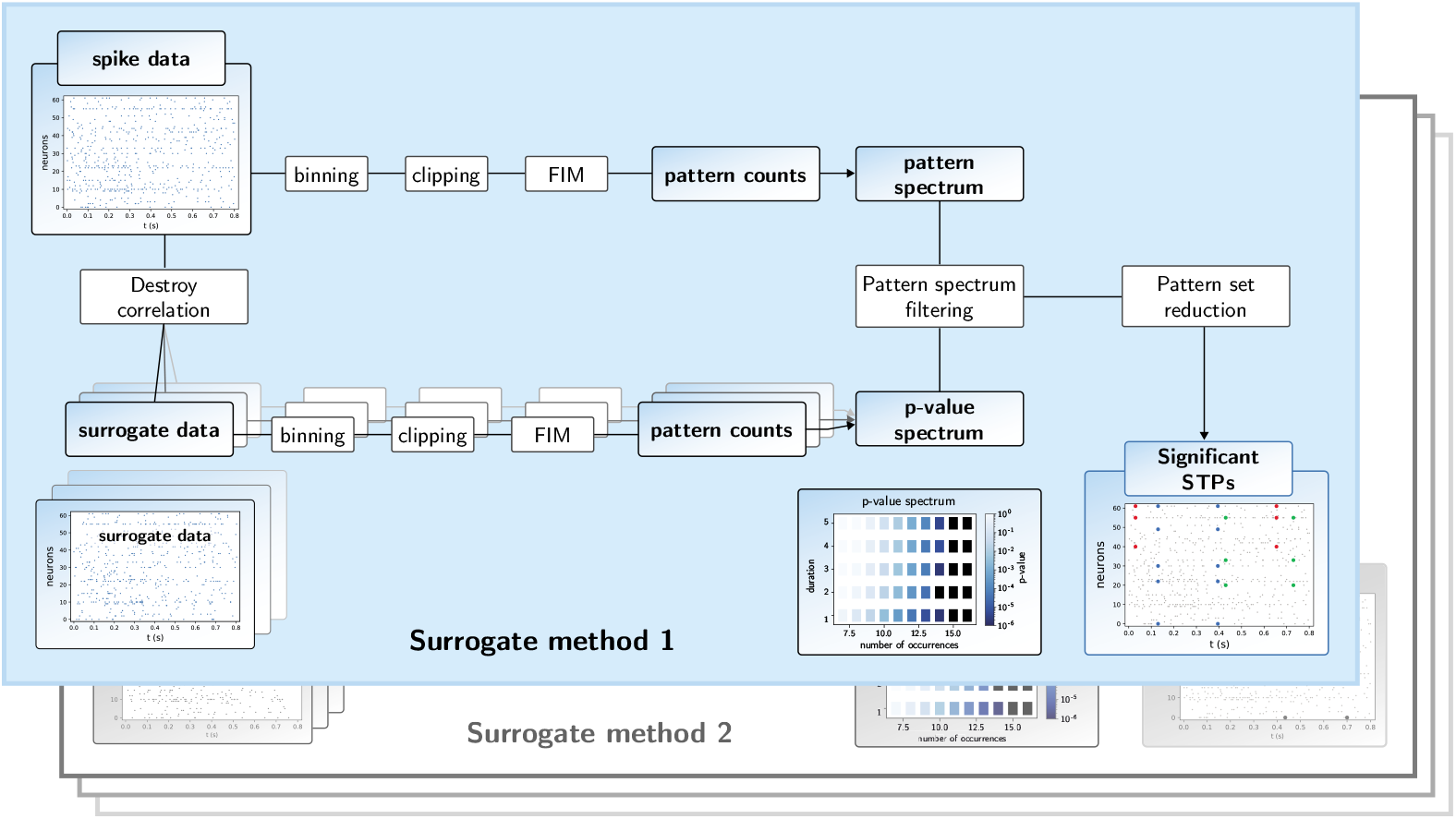
Workflow of the SPADE analysis. The top branch of the SPADE workflow shows the sequence of analysis steps of the original data until the pattern spectrum is derived. The bottom branch of the workflow starts with the generation of the surrogate data from the original data, followed by the same analysis steps as for the original data. The multiple overlapping panels in the lower branch indicate that this surrogate procedure is repeated many times, by which the p-value spectrum is built up. This then serves for the extraction of significant patterns through pattern spectrum filtering. After the application of the pattern set reduction, the significant STPs are provided as a result. ’Surrogate method 2’ indicates that the part ’Destroy correlation’ and ’surrogate data’ are replaced by another surrogate method, but the other steps stay the same.

**Figure 3.**
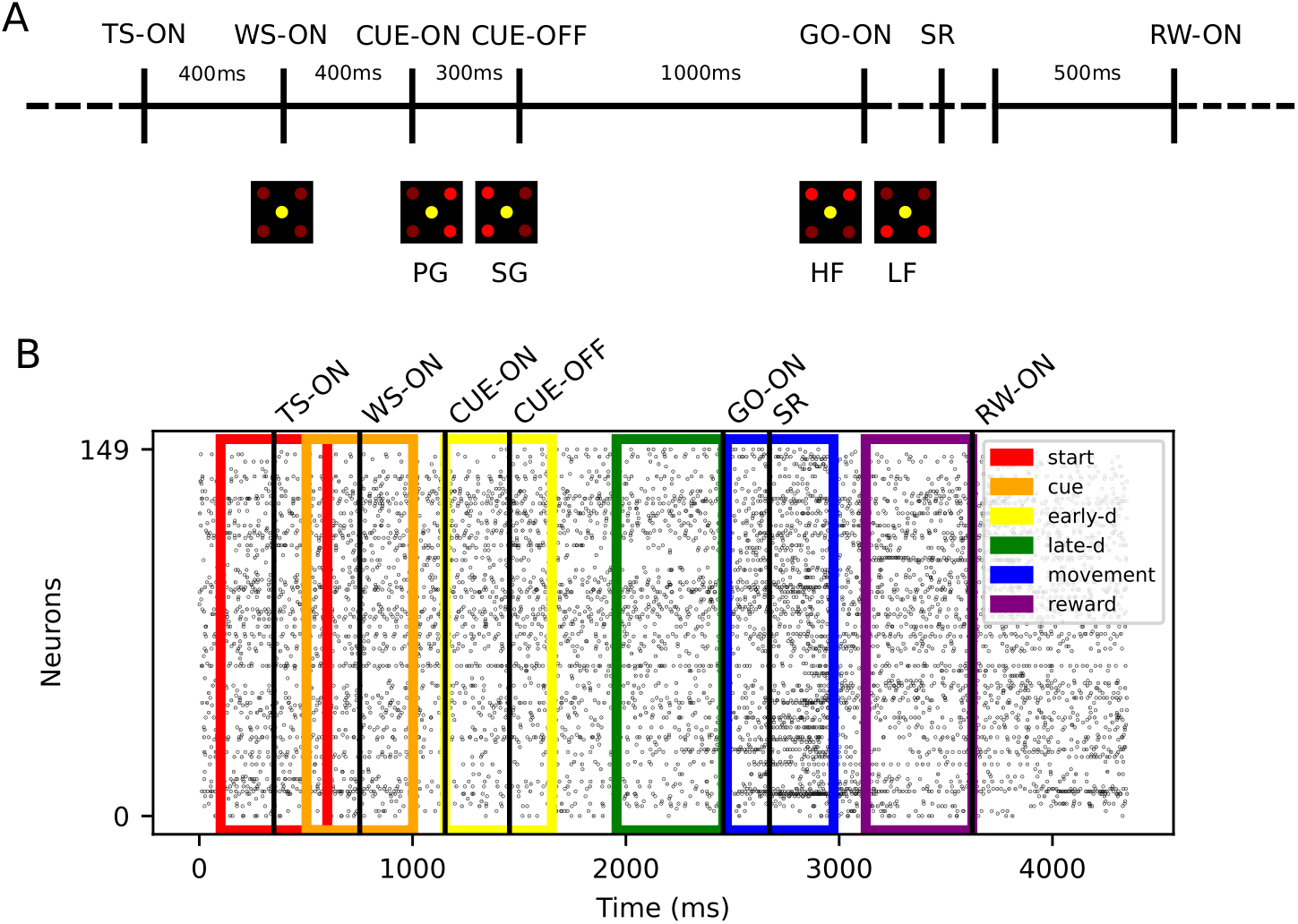
Experimental protocol and data preprocessing. (A). The trial start (TS) is self-initiated by the monkey. A waiting signal (WS-ON) prepares the monkey for the visual cue presented at CUE-ON, providing the grip type instruction (PG/SG). After 1000ms, the monkey is presented a second visual cue (GO-ON), specifying the force needed to pull the object (HF or LF) and the GO signal. The switch release (SR) marks the beginning of the movement. The monkey touches the object and maintains the grip for 500ms until the reward (RW-ON). The timing of the behavioral events SR and RW are variable and depend reaction time and movement speed. (B). The panel shows the simultaneous spiking activity of all neurons (*y*-axis) over time (*x*-axis) for one example trial (first successful trial of session i140703-001) of trial type PGHF. Each dot indicates one spike. The trials are aligned to trial start (TS-ON). The six colors rectangles represent the position of the six trial epochs (see legend).

**Figure 4.**
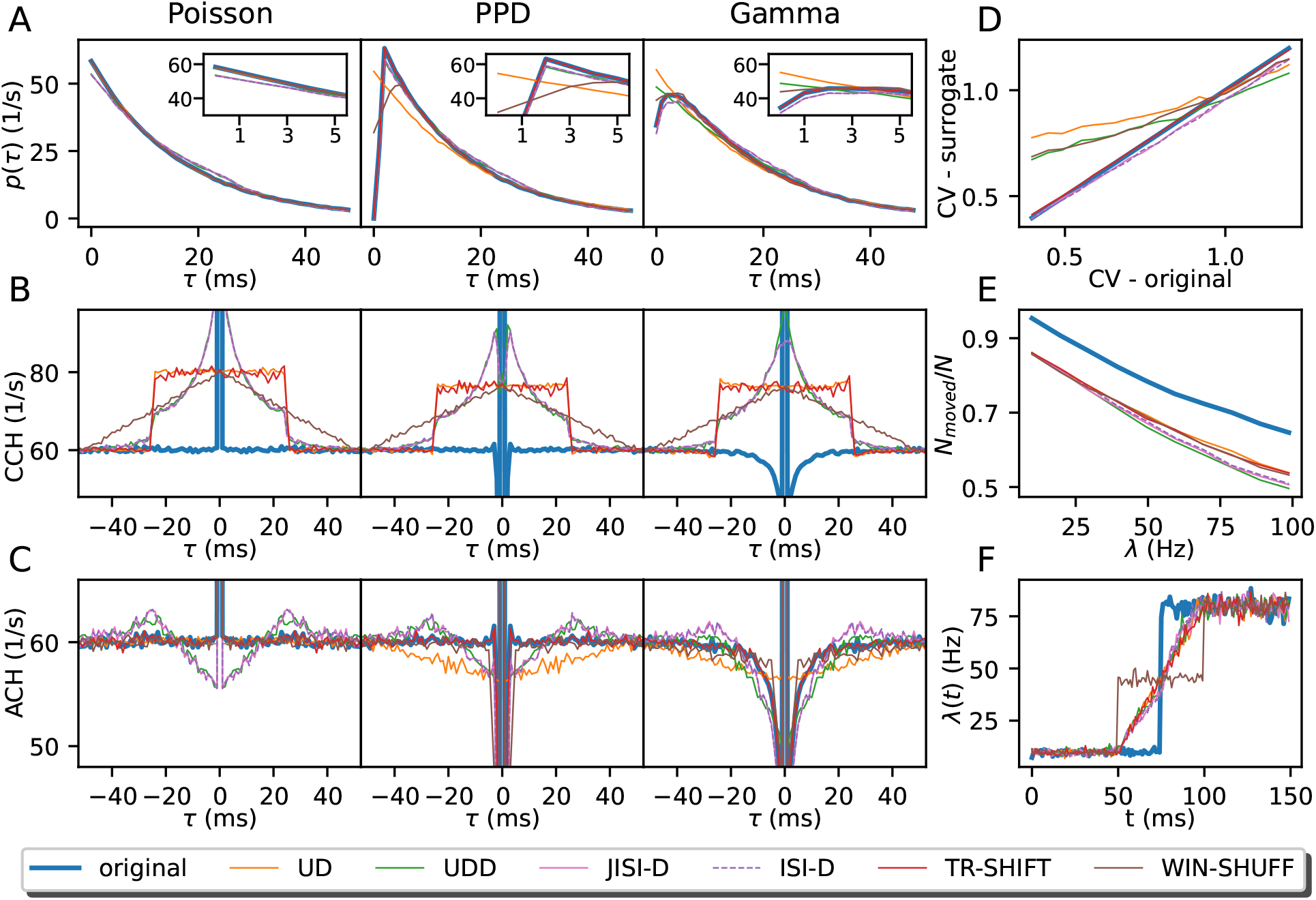
Overview of surrogate statistics. (A). The ISI distributions of original and surrogate spike trains (UD: orange, UDD: green, JISI-D: pink, ISI-D: violet, TR-SHIFT: red, WIN-SHUFF: brown) are shown as a function of the time lag *τ* in milliseconds (resolution of 1ms). For each process, the corresponding spike trains have a firing rate of 60Hz and an average spike count of 500, 000 spikes. The ISI region smaller than 5ms, is shown in an inset at the upper right corner. (B). The panel shows the cross-correlation between the original spike train (Poisson, PPD, and Gamma, in left, middle, and right column, respectively) with each of the surrogates (same color code as in A), blue is the correlation with the original spike train with itself (i.e., the auto-correlation) as reference. (C). Auto-correlation histograms are shown before (solid blue) and after surrogate generation (colored lines). For B and C, the x-axis shows the time lag *τ* between the reference spikes and the surrogate spikes (B) and the other spikes in the spike train (C). For panels B and C, we use the same data as in panel A. In panels D, E, and F, we only examine Gamma spike trains. (D). The panel displays the relation of the original CV (x-axis) against the CV of the surrogates (y-axis). Parameters are the same as in Panels A, B, and C (right), but we vary the CV 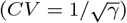 in steps of 0.05, ranging from 0.4 to 1.2 (*λ* = 60Hz). (E). The ratio of moved spikes (*N*_moved_) over the spike count *N* is shown. We show it as a function of the firing rate from 10 to 100Hz in steps of 10Hz on the x-axis (*γ* = 1.23). (F). In the panel, we show the conservation of the rate profile of a Gamma spike train (*γ* = 1.23) and its corresponding surrogates. The firing rate change is a step function, going from 10Hz to 80Hz (10, 000 realizations, spike train duration of 150ms), and is computed as a PSTH with a bin size of 1ms.

**Figure 5.**
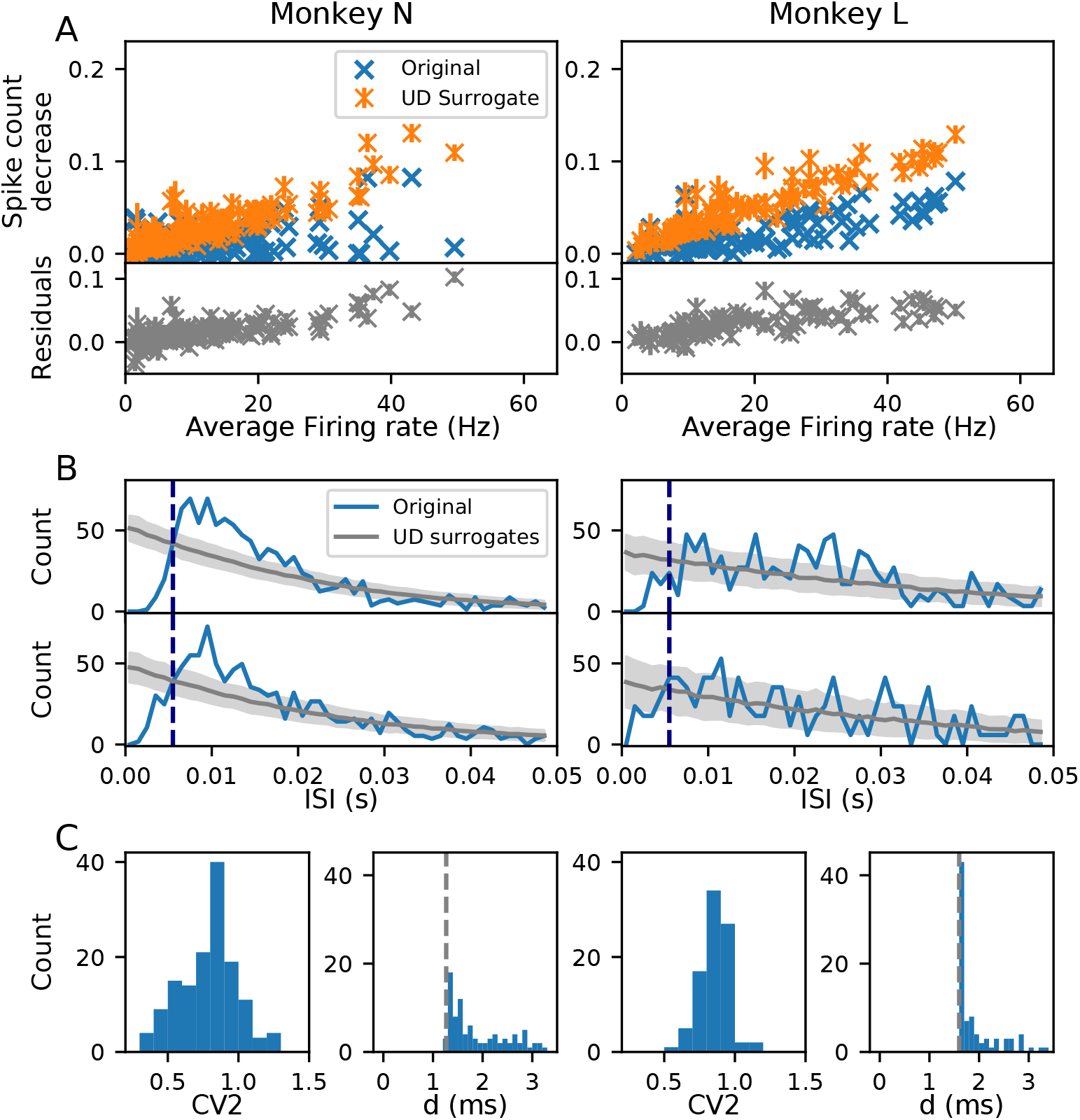
Modification of spike trains due to binarization. (A). Spike count reduction resulting from binarization and UD surrogate generation. Results from the analysis of two experimental data sets (sessions i140703-001 and l101210-001) in the movement epoch of the trial type precision grip-high force (PGHF) of monkeys N (left) and L (right). Top panel: Spike count decrease as a function of the average firing rate. Blue crosses indicate the spike count reduction caused only by binarization of the original spike trains (one cross per neuron). Orange marks show the spike count reduction after surrogate generation by UD and binarization. The spike count reduction is normalized by the spike counts of the original continuous-time spike train. Orange bars indicate the standard deviation of the spike count reduction calculated across 100 surrogates. Bottom panel: residuals (gray) computed as the spike count difference between the original binarized spike trains (blue) and the UD binarized surrogates (orange), normalized as in the top panel. (B) and (C). Interval statistics of the data. B shows the ISI distribution of 2 neurons from monkey N (left) and 2 neurons of monkey L (right; in blue). Neurons represented are 7.1 and 16.1 for monkey N, 5.1 and 6.1 for monkey L, with channel-id.unit-id notation. In gray are the ISIs of the respective UD surrogate distributions with the mean (dark gray) and the standard deviation of 500 surrogates in light gray. The bin size of the binning (here: 5ms) is shown by the dashed dark blue line. In C the CV2 distributions are shown for all neurons in each of the data sets (C, left subpanel). C, right subpanels, show the respective minimal ISI from the ISI distributions of all neurons. The dead-times assigned by the spike sorting algorithm are indicated by the dotted gray line (1.2ms for monkey N and 1.6ms for monkey L).

**Figure 6.**
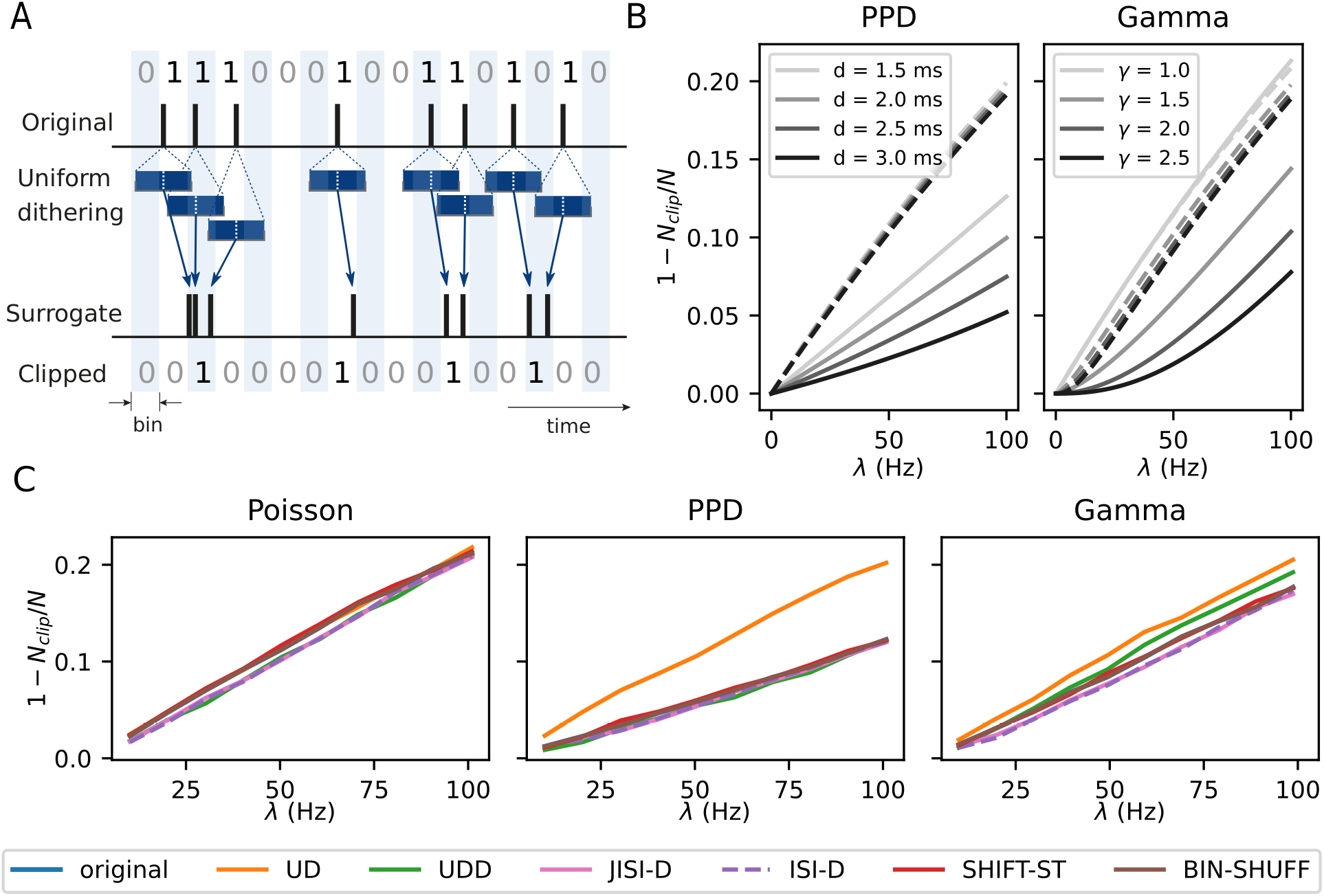
Origins of spike count reduction. (A). The sketch shows how a regular spike train is binarized. Below, it is illustrated how UD may change the spike times such that multiple spikes end up in single bins. The resulting binarized surrogate spike data are shown at the bottom. In contrast, due to the regular ISIs of the original process, its binned data are hardly losing spikes in comparison to the dithered version. (B). Analytical derivation of spike count reduction (after binning in 5ms intervals and clipping) for renewal point process models (PPD, left and Gamma process, right; solid lines, respectively), each with 4 different parameter sets (PPD: *d* = 1.5, 2.0, 2.5, 3.0ms, Gamma: *γ* = 1, 1.5, 2>, 2.5, different gray shades). The dashed lines show the same quantity for their UD surrogates. The firing rate of the processes is also varied and shown along the x-axis. The spike count reduction is shown on the y-axis, expressed as 1 *− N_clip_/N,* where *N_clip_* is the number of clipped spikes and *N* is the total number of spikes. (C). Spike count reduction of artificially generated spike train data ofFig 4 after binarization (bin width of 5ms) together with the corresponding surrogates in different colors. The firing rate is constant for each spike train and varies across realizations from 10 to 100Hz in steps of 10Hz (along the x-axis). The spike train durations are fixed such that, given the firing rate, all spike trains have an expected spike count of 10, 000 spikes.

**Figure 7.**
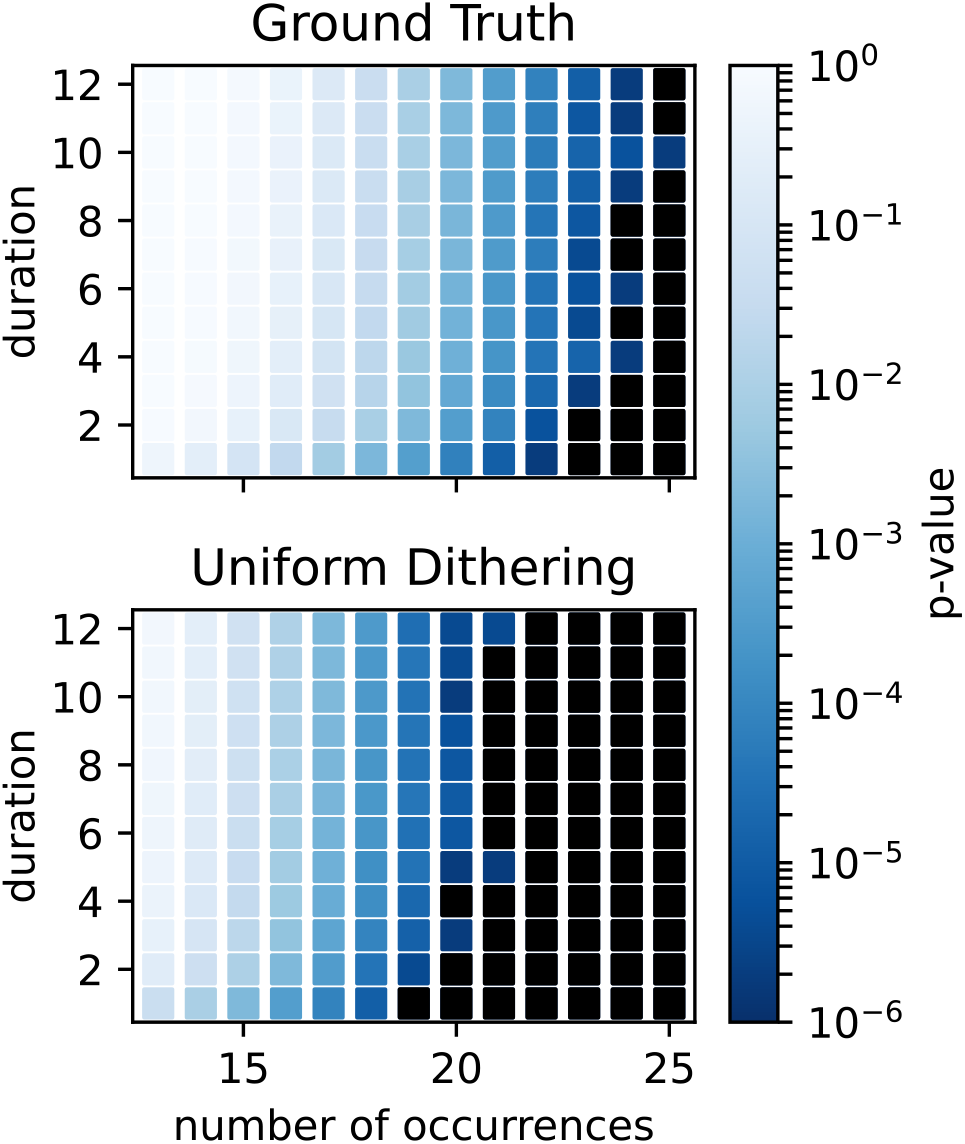
Consequences of spike count reduction.P-value spectrum of the original data (top) and of their UD surrogates (bottom) for mined patterns (of size 3 only) for a range of different pattern durations d (y-axis), and pattern counts (x-axis). The p-values are expressed by colors ranging from dark blue to light blue (see the color bar, identical for both spectra). The bin size is 5ms. The PPD data are of 100 realizations of *n* = 20 parallel independent spike trains with parameters *λ* = 60Hz, *d* = 1.6ms. The p-value spectrum for its uniformly dithered surrogates of the PPD data are derived from 5000 surrogates, dither parameter Δ = 25ms.

**Figure 8.**
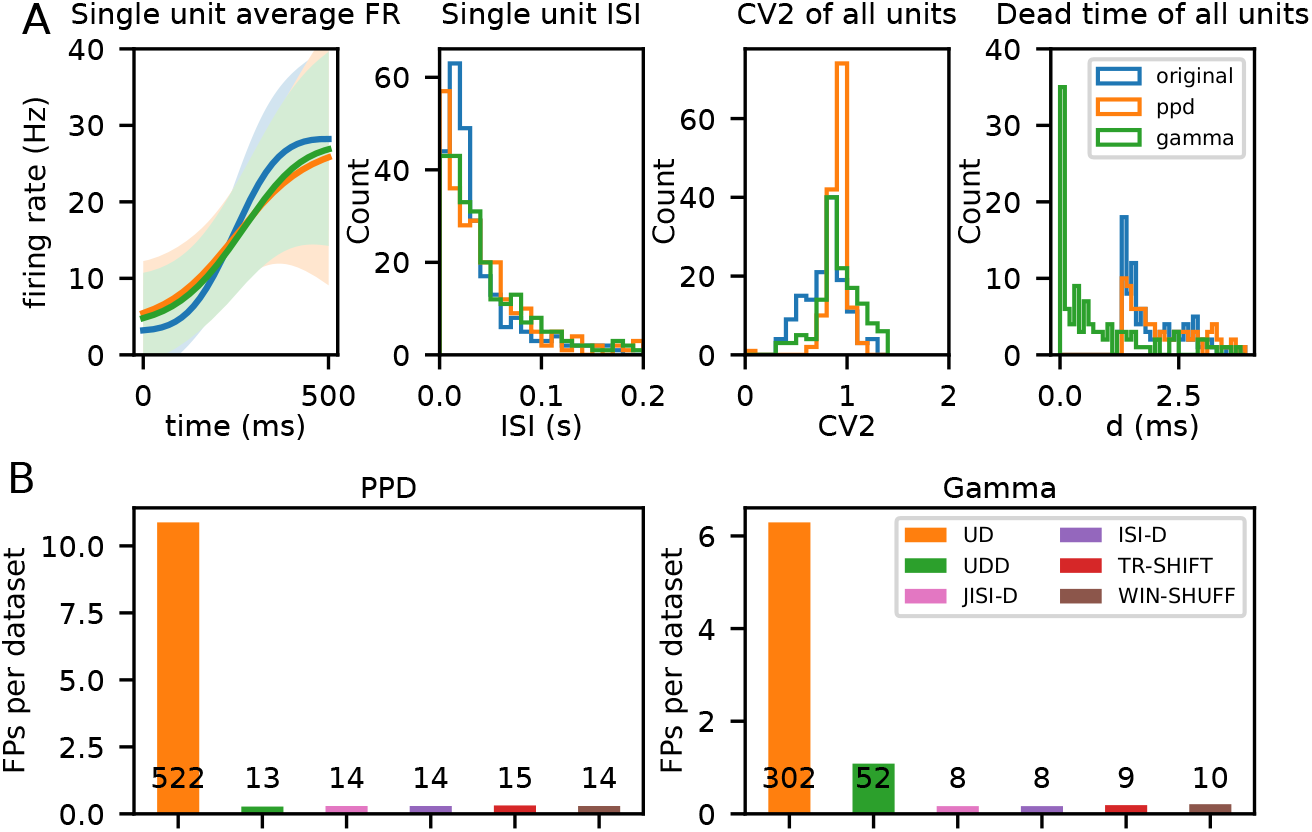
Evaluation and analysis of false positives across surrogate techniques for pattern detection with SPADE. (A). Comparison of statistics of the original to the generated artificial data during the movement epoch (PGHF trial type). In blue, orange and green we represent original, PPD and Gamma data spike trains respectively. Left graph: average firing rate of a single unit across one epoch of 500ms; second from left: ISI distribution of a single unit; third from left: average CV2 estimated trial wise for all neurons; fourth from left: dead-time as minimum ISI for all neurons. (B). number of false positives (FPs) detected across surrogate techniques (color-coded) normalized over the 48 (2 sessions *×* 6 epochs *×* 4 trial types) data sets analyzed, left for PPD and right for Gamma process data analyses. Numbers in text represent the total number of FPs over all data sets per surrogate technique. (C) and (D). Average firing rate of neurons for each monkey (N in panel C and L in panel D), epoch (y axis) and behavioral type (for each epoch ordered as PGHF, PGLF, SGHF, SGLF). Left PPD data, and right Gamma data. Colored dots represent individual units involved in FPs: blue dots indicate the average firing rate of units involved in FP patterns found for all surrogate techniques, orange dots for UD surrogates, green dots for UD and UDD only, and red dots for other combinations of different surrogate techniques. Grey dots represent the average firing rate of individual units not involved in any FP.

**Figure 9.**
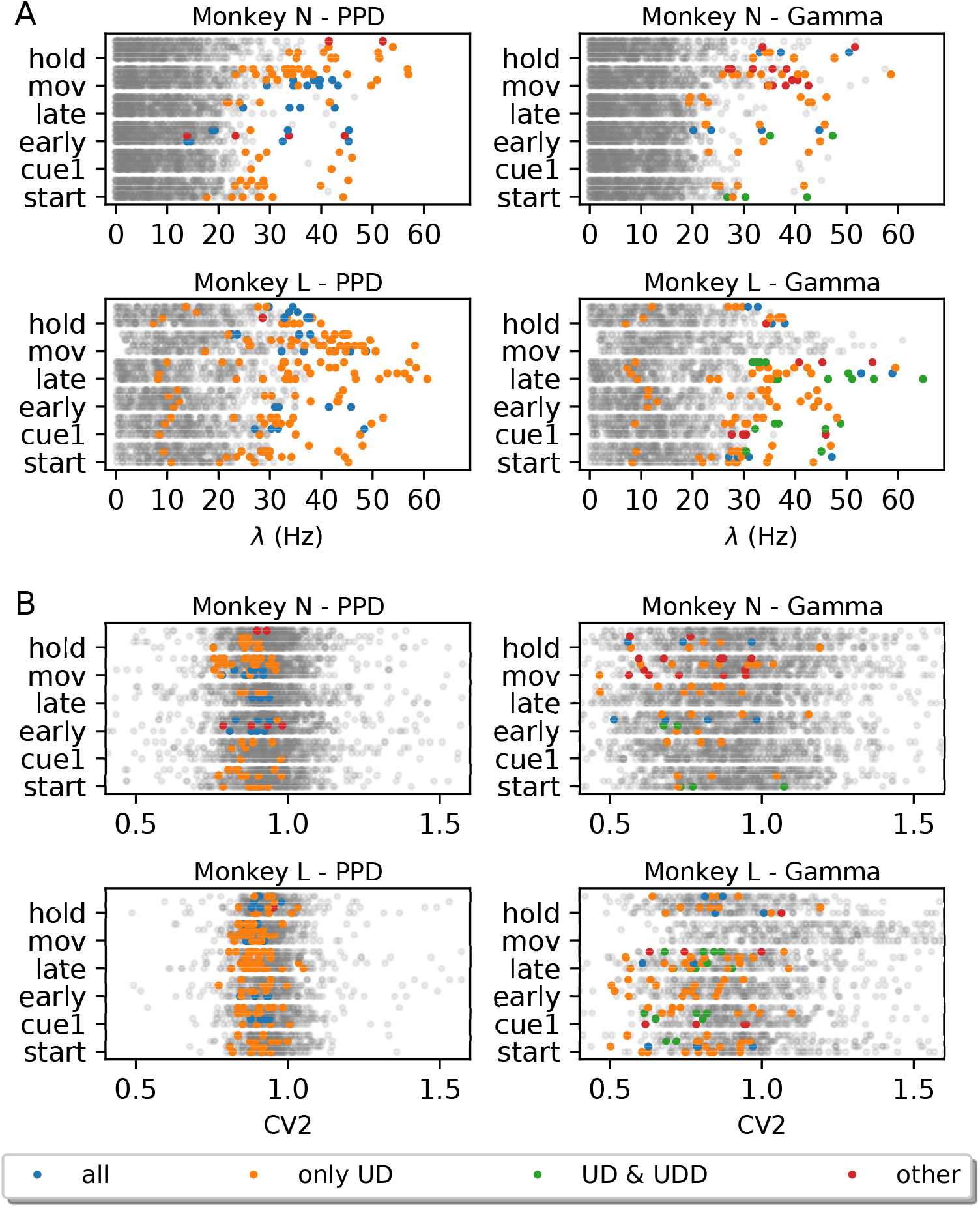
Average Firing rate and CV2 of neurons participating in FP patterns against all neurons. (A). Average firing rate of neurons for each monkey (N at the top and L at the bottom), epoch (y-axis) and behavioral type (for each epoch ordered as PGHF, PGLF, SGHF, SGLF). Left PPD data, and right Gamma data. Colored dots represent individual units involved in FPs: blue dots indicate the average firing rate of units involved in FP patterns found for all surrogate techniques, orange dots for UD surrogates, green dots for UD and UDD only, and red dots for other combinations of different surrogate techniques. Grey dots represent the average firing rate of individual units not involved in any FP. (B). Average CV2 of neurons for each monkey (N at the top and L at the bottom), same structure as (A).

**Figure 10.**
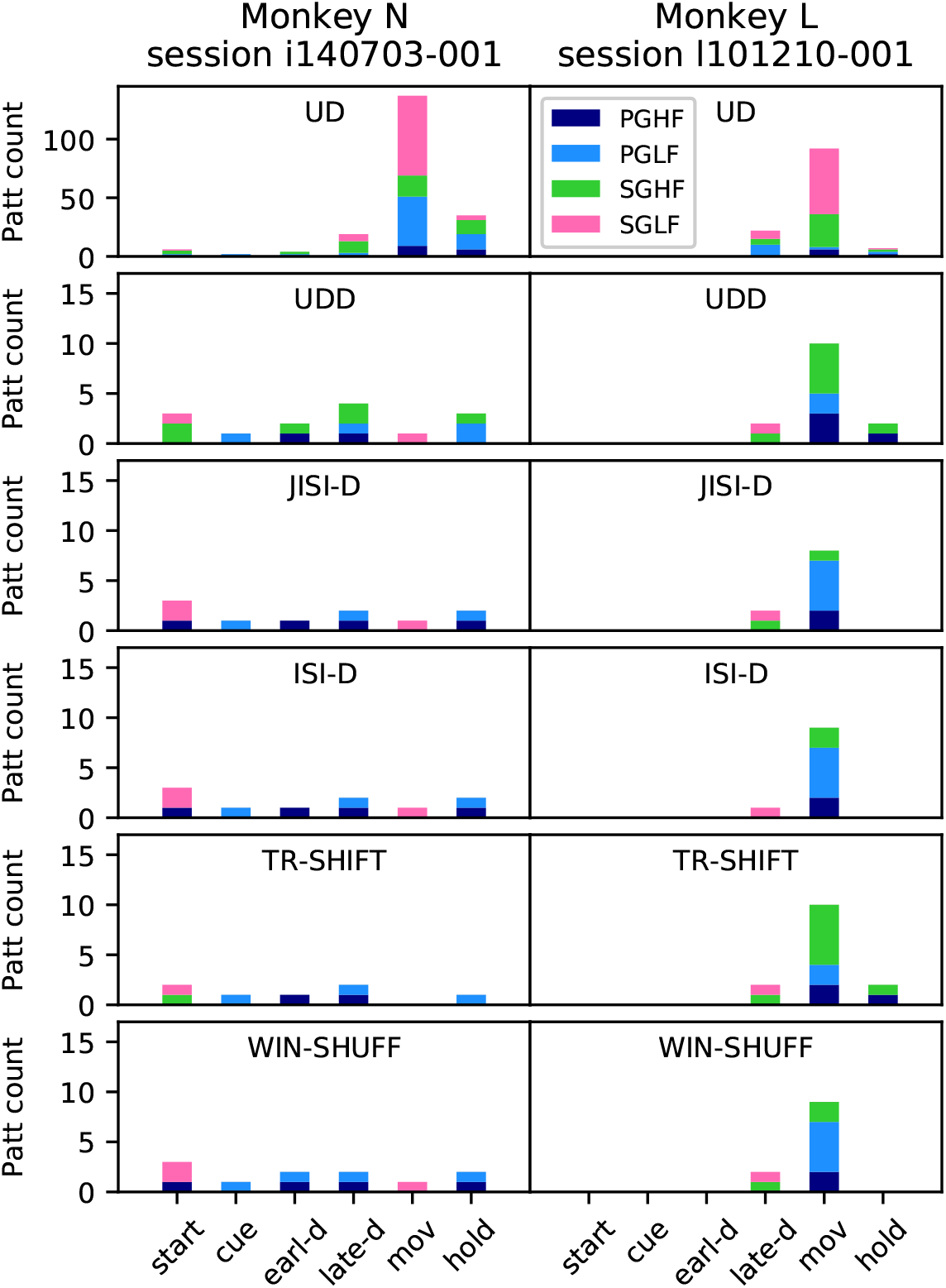
Analysis of experimental data. SPADE results for two sessions of experimental data: session `i140703-001` (left) and session l101210-001 (right). Histograms represent the number of significant patterns detected by SPADE in each epoch (start, cue, early-delay, late-delay, movement and hold), color-coded according to the grip type (precision/side grip -PG/SG- and low/high force -LF/HF). Each row corresponds to one surrogate technique (note the different y-axis scale for UD).

## 3 Results

### 3.1 Statistical comparison of surrogate methods

To get a better understanding of the effects of the surrogate methods on the statistical features of the spike trains, we first perform a comparison on stationary and independent data. For this purpose, we simulate point process models with well-defined properties: a Poisson process as a reference, a Poisson process with dead-time (PPD), and a Gamma process. The latter two are chosen to mimic the ISI distributions of the experimental data (Fig 5B). Here, the processes are stationary to exclude yet another statistical aspect. We explore the effect of all six surrogate methods on the statistical properties of the stationary and independent data (’original data’). The parameters of the data models are adjusted to be close to the experimental data and thus enable the transfer to experimental data to be analyzed later (section 3.4). In the next section (3.3), we will include non-stationary firing rates.

Fig 4 summarizes the results on stationary and independent data. The columns refer to the different spike train models (Poisson, PPD, and Gamma, from left to right, respectively) and in A the ISI distributions, in B the cross-correlation between the original and the surrogate data, and in C the auto-correlation of the original and the respective surrogate data for comparison. The 4th column displays a comparison of the CV of the original vs the surrogate data, the effectiveness of the displacement of the spikes through the surrogates, and the change of a rate step through the surrogate (from top to bottom).

#### 3.1.1 ISI distribution

The ISI distributions (Fig 4A, left; shown for *λ* = 60Hz) indicate for all surrogate data an exponential decay, lower for short ISIs for JISI-D, ISI-D, and UDD. Moreover, for Poisson data and in particular at high rates, the ISIs are often short. For the PPD process, UD dithers many spikes into the interval of up to 5ms (see inset), corresponding to the bin width. The other surrogates, which preserve the dead-time, follow more closely the original ISI distribution. On the other hand, a Gamma process does not contain a strict dead-time, but has a preferred ISI defined by the order of the process if *γ* > 1 (here *γ* = 1.23). The ISI distributions (Fig 4A, right) are most changed for UD and UDD, whereas the other methods maintain the ISI distribution at small ISIs almost identical to the original one.

#### 3.1.2 Similarity of original and surrogate data

As a second step, we take into consideration the cross correlation between different surrogate realizations and the original process. For the Poisson process (right), the different surrogates seem to generate again a Poisson-like process, since all surrogates move spikes around the original spike positions (Fig 4B, left): uniformly within the dither window for UD or TR-SHIFT, or in a triangular fashion for WIN-SHUFF, high probability around zero and exponential-like decay for JISI-D, ISI-D, and UDD up to the dither window of 25ms. In case of JISI-D, ISI-D and UDD the dithering is limited by construction to the interval between the former and the next spike. Thus, for Poisson data and in particular at high rates, the available dither window may not even be used completely, but only within the time interval between the two limiting spikes. Therefore, spikes are on average less moved and stay relatively close to their former positions.

For the case of PPD, the distribution of the spike shift from its original position (Fig 4B, middle) is similar to the Poisson process case (same panel - left), but the probability that spikes stay close to their original position is slightly reduced (e.g., pink and dashed violet and green have a lower peak close to the center). Finally, for the Gamma process, spikes are shifted into short ISIs, and to a lesser degree for WIN-SHUFF. The shift of the surrogate spikes from the original data (Fig 4B, right) is similar to the case of the Poisson and PPD models.

#### 3.1.3 Auto-correlation

For the Poisson process, the auto-correlations (Fig 4C, left) for UD, TR-SHIFT and WIN-SHUFF are flat besides the center peak, whereas JISI-D, ISI-D, and UDD show a decreased probability for very small ISIs, and then an increase up to the maximum dither width Δ = 25ms with a peak above baseline. The reason for this difference is the limitation that spikes may not be exchanged in their order, as for the other methods. For example, if the reference spike is close to the following and not to the preceding one, it will be more likely displaced backwards in time than forwards. This is also slightly visible in a difference of the ISI distributions (Fig 4A, left), compared to the other methods. Moreover, the increase towards Δ above baseline is due to the shift of dither probability to higher time intervals. When looking at the PPD process, we notice instead that for JISI-D, ISI-D, and UDD, the auto-correlations show - as compared to the corresponding surrogates of the Poisson process - also reduced short ISIs, but to a lesser extent. UD moves spikes into small ISIs, but on a smaller scale than the Poisson process (Fig 4A, left and middle, orange), and therefore shows a reduced probability when two spikes have a time difference of less than twice the dither parameter (here 50ms). Finally, in the Gamma process we have seen that the ISI distribution is maintained at small ISIs almost identical to the original one. Similarly, are the auto-correlations (Fig 4D, right): TR-SHIFT is identical to the original process; JISI-D and ISI-D are mostly preserving the auto-correlations, but still have a small bump above baseline at around Δ due to the dither restriction not to dither beyond the former and the next spike. WIN-SHUFF has a sharp reduction of spikes after the reference spike, and UD has a dip around 0.

#### 3.1.4 Coefficient of Variation of ISIs

We learned that the ISI distributions are affected by most of the surrogates. Fig 4D illustrates how the coefficient of variation (CV) of the surrogates differs in contrast to the original Gamma process (rate fixed to 60Hz), where the CV ranges from 0.4 to 1.2 in steps of 0.05. Non-preservation of the CV in the surrogate data as compared to the original data can be a potential source of false positives, in particular for very small CVs or CVs > 1 (Pipa et al., 2013). To facilitate the comparison, we also show the diagonal (blue). UD changes the CV the most, from original 0.4 to 0.75, i.e., losing strongly its high regularity, and increases even more, with a low slope, to a maximum slightly over 1.0 for the original CV of 1.25, so here burstiness is reduced. WIN-SHUFF and UDD behave similarly to UD, but for CV= 0.4 of the original data, these surrogates have a lower CV than UD (Fig 4D; orange, green, and brown lines above all others); moreover, UDD stays below UD for all CVs. WIN-SHUFF has a slightly higher slope and reaches a maximum still below the original Gamma process.

JISI-D, ISI-D and TR-SHIFT start with identical CVs than Gamma, and TR-SHIFT keeps the same CV as the Gamma CV for all CVs. However, JISI-D and ISI-D have a lower slope than the Gamma process, but still reach high values about 0.05 less than the highest CV at 1.25.

In summary, although the ISI distributions seem not to be strongly affected, the effect on the CVs can be very strong. For UD, UDD, and WIN-SHUFF, the CV slightly changes (in both directions), and for JISI-D and ISI-D, the CV decreases. A strong change in the CV of the surrogates can lead to false positives (Pipa et al., 2013). Only for TR-SHIFT, the CV is unchanged.

#### 3.1.5 Ratio of moved spikes

Next, we study if spikes are moved from their original bin in their surrogate realization. The reason of this interest is that if spikes are not sufficiently moved, correlations are not destroyed as intended, and thus may lead to false negatives. Therefore, we measure the ratio of the number of spikes that are displaced from their original bin position relative to the total number of spikes. We generate original data and its surrogate data and vary the firing rate (from 10 to 100Hz in steps of 10Hz, Fig 4E). If two spikes exchange their bin positions they are both considered as not moved. The spike ratio is also shown as a reference for two independent realizations of a Gamma spike train (*γ* = 1.23, blue line).

With increasing firing rates, the ratio of moved spikes decreases for the surrogates. Ideally, the surrogates should be similar to the effect attained on the original process, i.e., the colored lines in Fig 4E, should be as close as possible to the blue line. However, none of the surrogate techniques meets this ideal setting, and there are constantly 10% fewer spikes moved as compared to the blue line, i.e., the ratio of moved spikes for two independent spike train realizations. Nonetheless, we observe for all surrogates that the ratio of moved spikes decreases with increasing firing rates, which corresponds to the fact that increasingly more bins are already occupied and thus the resulting binned surrogate spike train is more similar to the binned original. UDD, ISI-D, and JISI-D displace fewer spikes, in particular for higher firing rates as compared to UD, TR-SHIFT, and WIN-SHUFF. The fewer the spikes that are not effectively displaced, the higher the peak at zero-delay of the cross-correlation of the original and the surrogate data (Fig 4B). Almost 50% of JISI-D, ISI-D, and UDD are not moved at 100Hz, and, for lower rates, they remain below the ratios of WIN-SHUFF, UD, and TR-SHIFT. As a consequence, we can expect that JISI-D, ISI-D, and UDD in general tend to yield more false negatives than WIN-SHUFF, UD, and TR-SHIFT.

#### 3.1.6 Rate change in surrogates

Changes in the firing rate profile of the surrogates compared to the original data may be a source for false positives (Grün, 2009). An optimal surrogate method should follow as closely as possible the original firing rate profile. Therefore, we test here an extreme case where the original data have a rate step (as in Louis et al., 2010a), jumping from 10Hz to 80Hz (Fig 1F). We observe that for all surrogates but WIN-SHUFF the firing rate step is converted into a linear increase, which starts at 25ms (dither parameter Δ) before the step and ends at 25ms after the rate step. This behavior has already been derived analytically and observed in Louis et al. (2010a) for UD: it corresponds to the convolution of the original firing rate profile with the dither (boxcar) function. WIN-SHUFF introduces a second step in the firing rate profile, which is due to the fact that it generates a locally-stationary firing rate within the shuffling window (here 50ms). We conclude that all surrogate techniques smooth the original firing rate profile, whereas WIN-SHUFF creates an additional intermediate rate step. Thus, for steep increases in the firing rate profiles, we have to expect the occurrence of false-positive patterns due to this smoothing.

#### 3.1.7 Summary of the effects on the spike-train statistics of surrogates

We explored different aspects of the statistics of the surrogate spike data as compared to its original process. In general, surrogate data are not identical to the original data, but change to a different degree. The effects for the three data models are summarized and not differentiated, since they anyway are similar. These are listed in Table 1. Features that are preserved are indicated by ’yes’, approximately preserved (’approx.’), and not preserved (’no’).

**Table 1.** Table summarizing the statistical properties of the six discussed surrogate techniques conserve (yes/ approx./ no). The dead-time conservation is evaluated based only on results of PPD process, otherwise on the results for all data models (Poisson, PPD, Gamma).

| Feature/Method | UD | UDD | ISI-D | JISI-D | TR-SHIFT | WIN-SHUFF |
| --- | --- | --- | --- | --- | --- | --- |
| ISI | no | no | approx. | approx. | yes | approx. |
| dead-time | no | yes | yes | yes | yes | no |
| Auto-correlation | no | no | no | no | yes | approx. |
| Firing rate modulation | approx. | approx. | approx. | approx. | approx. | approx. |
| Spike train regularity (CV < 1; regular) | no | no | approx. | approx. | yes | no |
| Spike train regularity (CV > 1; bursty) | no | no | approx. | approx. | yes | approx. |

### 3.2 Impact on spike counts after spike train binarization

A typical step in the analysis of experimental spike data is to downsample them to the time scale of relevance, e.g. ms resolution. This is typically done by binning the continuous spike trains to bins of a few ms width, resulting in spike counts per bin. Further analysis steps often require to reduce these data to 0-1 sequences (e.g., as described for the SPADE method in sec. 2), thus the bin contents is reduced to 1 if one or more spike are in a bin (’clipping’), or to 0 for no spike. Thus, we now explore if and how the binarization step affects surrogate data. For doing so, we compare the spike counts per neuron before and after the binarization step, for both the original and surrogate data. We notice that for some neurons the total spike counts of the UD surrogates are much lower than those of the original data. Further analysis of this aspect shows that the higher the firing rate of a neuron the larger the spike count reduction and thus the corresponding mismatch. Fig 5A shows that for two different data sets, each from a different monkey, and we find a spike count mismatch of up to 10% between the UD surrogates and the original data (Fig 5A, bottom, gray). Such a difference in the spike count is troublesome, since it is expected to lead to a reduced pattern count of the surrogates as compared to the original data and, thus, is expected to yield an overestimation of the significance of patterns.

#### 3.2.1 Origin of spike count reduction

The UD procedure as such is not deleting any spike, only the binarization step does. In this regard, the latter step is crucial: when applied to the original and the surrogate data, it leads to different spike counts. Here we aim to understand why this is the case. One potential reason is the change of the inter-spike interval (ISI) statistics of the spike trains with and without dithering, as already analyzed for stationary spike models in sec. 3.1.1. Fig 5B, shows the ISI distribution for two example neurons of the experimental data (in blue; right for monkey N, left for monkey L) and for comparison, the ISI distributions of the uniform dithered surrogates (gray). In the experimental data, the ISI distribution is peaked at a certain ISI, here between 5 and 10ms, but the ISI distributions of the surrogate data are decaying exponentially, and thus also fill small ISIs. This indicates the fact that the original spike trains are more regular than the surrogates and so small ISIs have a lower probability. The regularity of the experimental data is confirmed by the measurements of the CV2. Indeed, the CV2 distribution of all neurons of each of the two data sets (Fig 5C, left subpanels, in both columns) is rather below 1, i.e., more regular than Poisson.

In addition, the distributions of the minimal ISI of each neuron per data set (Fig 5C, right subpanels of the two columns) exhibit a minimal ISI of 1.3ms for monkey N and 1.6ms for monkey L. This corresponds to the dead-times of the spike sorting algorithm that cannot resolve overlapping spikes. The different dead-times for the two monkeys are due to a different number of sample data points considered for spike sorting (Brochier et al., 2018). The corresponding ISI distributions of the surrogate data (Fig 5B) show that there are ISIs smaller than the minimum ISI of the respective experimental data. Thus, the dithering procedure generates shorter ISIs in the UD surrogates than in the original data.

To verify our interpretation that the combination of UD and binarization causes the spike count reduction, we perform a similar analysis on well-defined ground truth data, i.e., simulated PPD and Gamma spike trains. For simplicity, we now choose both of a constant rate but adapt the dead-times or shape factors, respectively, to account for the ISI features of the experimental data.

Fig 6C shows the spike count reduction (expressed as 1 − *N_clip_/N*, where *N_clip_* is the number of clipped spikes and *N* the total number of spikes) that results after binning (5ms) of the PPD (left) and Gamma process (right) with (dashed) and without (solid) application of uniform dithering. The analytical derivation can be found in Stella et al. (2021). The graphs show the spike count reduction as a function of the firing rate of the processes. The PPD model shows a higher spike loss the higher the firing rate, and a lower spike reduction for larger dead times. The uniformly dithered PPD show for all dead-times an increase in the spike reduction with higher firing rates, but to a larger extent than for the original PPD processes. The Gamma process (right) for *γ* > 1 shows a similar result as for the PPD: an increase of spike count reduction for increasing firing rates, and the larger the shape factor, the lower the spike count reduction. The increase is rather parabolical compared to the PPD. The Poisson process (light gray, *γ* = 1) shows a much larger and linear increase of spike reduction with rate, more strongly than for Gamma processes with *γ* > 1 (darker grays).

So, a) why does a Poisson-like process lose more spikes through binarization than a process with a non-exponential ISI distribution, and b) why does uniform dithering lead to a loss of spikes compared to the original experimental data? As shown above (Fig 5B), uniform dithering generates a more Poisson-like ISI distribution. Such a process contains spikes that follow each other in short intervals. Such a cluster of spikes may fall within a bin, and then the content of the bin is reduced to 1 by clipping (see for illustration Fig 6). A PPD process has a strict dead-time, leading to the fact that the probability of more than one spike in a bin is reduced, and thus there is a smaller spike loss. In fact, the goal of preserving the dead-time is to reduce the difference between the binned and clipped spike counts of the original spike train and its surrogates. The closer the dead-time to the bin size, the more unlikely it is that two spikes are dithered into the same bin (because of the dead-time constraint), and the spike counts of the surrogates are closer to the spike count of the original spike trains.

#### 3.2.2 Spike Count Reduction in different spike train models

Fig 6C illustrates the spike count reduction for all types of surrogates applied to the three different spike train models of section 3.1: the higher the rate the more spikes are lost. However, there are some differences in the degree of spike loss for different data models, which we now discuss separately.

For Poisson data, the spike count loss increases approximately linearly with the firing rate for all surrogates. This happens as well for the original Poisson process (blue), up to 20% for a rate of 100Hz. Only JISI-D, ISI-D, and UDD overall have a slightly lower loss. In the case of the PPD process the loss of spikes (Fig 6A, middle) is considerably reduced for the PPD compared to the other models. This is true also for its surrogates, besides UD (orange), which loses more spikes. This may be explained by the ISI distributions that we observed in Fig 4A: UD dithers many spikes into the interval of up to 5ms (see inset), corresponding to the bin width. For the Gamma process, the spike loss for the different surrogates (Fig 6C, right) is higher than for PPD, but lower than for Poisson. UD loses the most, JISI-D and ISI-D the least.

Only one method preserves completely the spike count as the original spike train, and it is WIN-SHUFF. In fact, this method conserves the number of spikes of the original data and the number of occupied bins in the original but already discretized data for each surrogate, i.e., there is no risk of spike count reduction (Table 1).

As UD surrogates evidence a strong spike count reduction in the context of binarization, we expect UD to yield a large number of FPs in a statistical test under this condition. An exemplary statistical test of this kind is the pattern spectrum filtering (PSF) used in SPADE. Further, UDD surrogates might lead to FPs in the case of regular data that do not exhibit a dead-time (e.g., Gamma spike trains). The study of the similarity of the surrogates to the original processes shows that JISI-D, ISI-D, and UDD might lead to fewer patterns detected, i.e., an underestimation of significance. Moreover, we expect FPs for WIN-SHUFF surrogates when the original firing rate profiles have a steep rate increase. Finally, the technique preserving the most statistics without showing any disadvantages is TR-SHIFT.

Before we move on, a note of caution concerning the use of the terms “false positives” and “false negatives” may be necessary: in the experimental data we don’t have an independent “ground truth” telling us which patterns are the “real” ones. So, in principle one cannot speak of false positives or false negatives, one can only speak about under- or over-estimation of significance probabilities. In the artificial data we can only generate data from a null-hypothesis, under which patterns only occur by chance. Therefore, any patterns found to be significant in these data are considered as false positives, and there can be no false negatives.

#### 3.2.3 Consequences of spike count reduction

Through the spike count reduction in the surrogate data also the expected occurrences of STPs may be reduced. This we prove by comparing independent data to UD surrogates. In Fig 7, we represent the results of such a comparison. We generate 20 parallel artificial independent PPD spike trains with a stationary firing rate of *λ* = 60Hz, and with a dead-time of *d* = 1.6ms, and a total length of 1s. From these spike trains, we generate 5000 surrogates, each with uniform dithering, discretize them and count the number of patterns using FIM. Based on the generated pattern spectrum, we calculate the corresponding p-values. In addition, we compare the p-value spectrum of the surrogate data (Fig 7, bottom panel) with the p-value spectrum of the ground truth data. To do so, we generate a large number of the independent PPD processes and analyze these by FIM to extract the patterns and generate the p-value spectrum. In other words, knowing the ground truth, we can use new spike train realizations as surrogates. Fig 7, top panel, shows the p-value spectrum of the PPD processes, for patterns of size 3, across different pattern durations. Since these data are independent, the resulting patterns occur by chance. The p-value spectrum of the surrogates of the PPD processes is shown below. The two p-value spectra illustrate that the p-values of the UD surrogates are smaller and UD surrogates have fewer pattern counts as compared to the ground truth data. In other words, pattern counts of the ground truth (PPD) data would become significant if compared to its UD surrogates although the ground truth data sets are independent. As a consequence, patterns in the independent original data would be classified as significant while being false positives.

From this analysis, we conclude that uniform dithering is not an appropriate surrogate method for spike data that either contain a hard dead-time or have a regular spiking behavior, as motor cortex data tend to have (Mochizuki et al., 2016). Therefore, we now deepen our evaluation of surrogate methods from section 3.1 by analyzing the impact of these on the SPADE analysis.

### 3.3 Performance of the surrogate methods

Next, we apply the six surrogate techniques to artificial data sets that are generated based on two experimental data sets (Brochier et al., 2018) to study the effect of the surrogate methods on the occurrence of false positives (Louis et al., 2010a). The experimental data are two sets of recordings from approximately 100 parallel spike trains from macaque monkey motor and pre-motor cortex during performance of a reach-to-grasp behavior (Riehle et al., 2013), explained in 2.3. We simulate non-stationary artificial data with the same firing rate profiles as the experimental data, and use as point process models a) the PPD to mimic the dead-time of the data (due to spike sorting), and also b) Gamma processes to account for their CVs. The spike trains are generated independently and hence all observed spike patterns occur by-chance and can be considered as false positives (FPs).

For the two experimental sessions, each of the 24 sets of concatenated data is modeled using the two point process models, resulting in a total of 2 24 2 = 96 data sets. The SPADE analysis is performed on all data sets, separately for each of the six surrogate techniques. We set the bin size to 5ms, the maximum pattern duration to 60ms, the significance level to *α* = 0.05, and use 5000 surrogates. For all surrogate techniques, we set the dither parameter to Δ = 25ms.

#### 3.3.1 False positive analysis

Fig 8B shows the number of retrieved false positive results for the PPD (left) and the Gamma data sets (right). For each point process, we show the number of FPs per data set and the total number of FPs (in text) across the six surrogate methods. Results show that we retrieve for all surrogate techniques, except for uniform dithering, a small number of FPs. For UD, we get for the PPD process in Fig 8B, left: UD (522; 88.1%), UDD (13; 2.2%), JISI-D (14; 2.3%), ISI-D (14; 2.3%), TR-Shift (15; 2.5%), and WIN-SHUFF (14; 2.4%). Contrarily, we see in Fig 8B (right) that the analysis of the Gamma data leads to the following number of FPs: UD (302; 77.6%), UDD (52; 13.4%), JISI-D (8; 2%), ISI-D (8; 2%), TR-Shift (9; 2.3%), and WIN-SHUFF (10; 2.6%). Considering all results, we conclude that UD leads to a very high number of false positives compared to the other surrogate methods, as we expected from our results of the previous sections. We also observe a relatively high number of FPs for the application of UDD on Gamma data, expected from our observations in section 3.2.2. The remaining surrogate techniques exhibit a similar number of FPs.

For further analyses, given these observations, we form four groups of FPs, depending on which surrogate techniques they are expressed in. The first and predominant group are the FPs present only in the SPADE analysis performed with UD surrogates. Second, we group together FPs present in all surrogate techniques. Third, in the case of Gamma data, we distinguish a subset of FPs found with both UD and UDD surrogates. Finally, we pool all FPs present in any other combination of surrogate analysis. To get an understanding of the rate properties of neurons that contribute to the FPs, we consider their average firing rates (over time and trials) and the group that they belong to (Fig 9A). In general, we find FPs in all analyzed data sets, but one (monkey L, movement, Gamma, all conditions). We observe that almost all neurons involved in FPs have an average firing rate higher than 20Hz. Neurons belonging to the first group are the largest set, and are present for both monkeys and data models, and in almost all data sets. The second group is present for both monkeys and models, but is more represented for PPD. The third group is present for both monkeys only for the Gamma model. This was already expected, given the higher spike count reduction, for UD and UDD in the case of Gamma spike trains (section 3.2.2). We also inspect the CV2, averaged over trials, of units involved in FPs (Fig 9A). FPs occur in neurons with a relatively low CV2 (0.7 *<* CV2 *<* 1), but this is not the case for neurons with very low CV2s (CV2 *<* 0.7; especially for monkey N). In almost all cases, bursty neurons (CV2 *>* 1) are not involved in FPs.

In summary, we observe that the surrogate technique leading to most FPs is UD, followed by UDD (only in the case of Gamma data). Neurons exhibiting an average firing rate higher than 20Hz, and having a CV2 *<* 1 are predominantly involved in FPs. Moreover, there is a small amount of FPs detected using all surrogate techniques, which is expected given a certain significance threshold. Nonetheless, regular and high-rate neurons are more prone to raise the false-positive rate (Harrison and Geman, 2009).

### 3.4 Application to experimental data

As the last step, we apply SPADE with the six surrogate techniques to the two sessions of experimental data introduced in section 2.3. Here our goal is to analyze with SPADE experimental data, for which we do not know the ground truth (i.e., the presence and amount of significant patterns) and show the differences resulting from the application of the surrogate techniques in the significance testing. In Fig 10, we present the found number of significant patterns for each epoch and trial type (different colors). The results are shown for each monkey separately, since their data differ in terms of CV2, dead-time and firing rates (Fig 8 and Fig 9). The exact number of patterns found is: UD (N:203, L:121), UDD (N:14, L:14), JISI-D (N:10, L:10), ISI-D (N:10, L:10), TR-SHIFT (N:7, L:14), WIN-SHUFF (N:11, L:11). Crucially, we detect more patterns (almost double the amount) in the analysis of experimental data than in the independently generated artificial data, except for UD and UDD (on Gamma data) surrogates.

A first observation is that the amount of significant patterns found using UD is much higher (note different y-axis scale) for both monkeys as compared to the other five surrogate techniques. Patterns occur mostly during the movement (mov) epoch where the firing rates are highest. Thus, given the calibration results from the former section, a large amount of those is likely to be FPs.

Taking from now on into consideration all surrogate techniques but UD, for monkey N (Fig 10, left column) we find patterns across all epochs, almost for all surrogates. The pattern numbers are relatively similar within and across epochs. Very specific is the pattern occurrence during the movement epoch (Fig 10, left column, pink): in fact, the same pattern occurs for SGLF behavioral context in all surrogates, but TR-SHIFT. During the start epoch, all surrogates show patterns in relation to SGLF, but some (UDD and TR-SHIFT) are also in relation to SGHF, and others (JISI-D, ISI-D and WIN-SHUFF) in relation to PGHF. In the cue epoch, all surrogates find patterns in relation to PGLF trials (Fig 10, left column, light blue). During early delay (earl-d) all surrogate techniques find patterns for PGHF trials, in addition, for UDD, a pattern for SGHF, and one in PGLF trials for WIN-SHUFF. During the late waiting epoch (late-d) patterns occur only in PGHF and PGLF trials (Fig 10, left column, blue and light blue), but we also observe patterns in SGHF trials for UDD (green, second row). Finally, during hold, we find patterns in PGLF trials for all surrogate techniques. In addition, we find a pattern in SGHF trials for UDD and a pattern in PGHF trials for JISI-D, ISI-D and WIN-SHUFF.

For monkey L (Fig 10, right column) most patterns occur during the movement epoch for PGHF, PGLF, and SGHF, however in slightly different combinations. During the late phase of the waiting period (late-d) four out of the five surrogates find the same patterns (1 for SGLF and 1 for SGHF). During the hold epoch only for UDD and TR-SHIFT we find the same patterns, 1 for PGHF and 1 for PGLF. We do not detect any significant patterns in the start, cue, and early delay epochs.

Interestingly, for both monkeys, the significant patterns are specific to the epochs, i.e., identical patterns do not repeat in different epochs, but the patterns are different in the temporal delays of the spikes and are mostly composed of different neurons. Previous studies on this experiment (Riehle et al. (2018), Figure 2) showed that monkey L has on average a shorter reaction time than monkey N, and has shorter and more pronounced rate increases within the movement epoch, which may explain the high amount of patterns during that epoch. In contrast, monkey N shows patterns already in the start epoch, and the number of detected patterns remains almost constant throughout epochs.

## 4 Discussion

The generation and use of surrogate data has become an important methodological approach to data analysis when the data are complex and there are no reasonable simple hypotheses that could be used for statistical evaluation. This holds in particular for complex experimental data, when the experiment has been repeated only a few times (i.e. small number of high-dimensional data points). Multiple spike train recording is a typical case.

Of course, it is still possible to use comparably simple spike-train distributions as null-hypotheses to test for significant repetitions of spatio-temporal spike patterns, which will find more significant patterns with much less effort (Staude et al., 2010; Russo and Durstewitz, 2017). However, there may be several potential reasons for a significant deviation from such a simple null-hypothesis, in particular when it assumes stationarity and does not include the obvious effects of covariation of neural firing rates with the experimental stimulation (Grün, 2009; Pipa et al., 2013).

In this study, we perform a comparison of six surrogate techniques which are used in the analysis of parallel spike trains.: uniform dithering (UD), uniform dithering with dead-time (UDD), window shuffling (WIN-SHUFF) -both newly introduced-, joint inter-spike interval dithering (JISI-D; Gerstein, 2004), inter-spike interval dithering (modified from Gerstein, 2004), and trial shifting (Pipa et al., 2008: Fig 1). Such surrogate approaches have the goal of destroying the exact spike timing relations between the neurons. We quantified which statistical features of the original spike trains are conserved in surrogates, examining the ISI distribution, the auto-correlation, the cross-correlation, the firing rate modulations, the ratio of moved spikes, and the coefficient of variation (Table 1). These were evaluated on stationary artificial data (Poisson, PPD, and Gamma spike trains) that contained relevant features of the experimental data (dead-time and regularity; Fig 4). Additionally, we observed that UD does not preserve the spike count when the spike train is binarized and leads to a very strong spike count reduction for the PPD model, not for the Poisson model, and less for the Gamma model.

The issue of spike count reduction shown for UD on stationary, independent data is particularly relevant, as it might influence a statistical test result. Looking more in detail into real experimental data, we have shown that the usage of uniform dithering (UD) as a surrogate technique, followed by binarization (binning and clipping) of the spike train, leads to a mismatch in the spike counts between the original and the surrogate data (Fig 5). The spike count reduction is worthy of attention and increases with the firing rate, which we verified analytically and through simulations (Fig 4A, Fig 6B). Moreover, we showed that two factors play a large role in the spike count reduction: the neuronal dead-time and the coefficient of variation (Fig 6B).

Evaluation of spatio-temporal patterns is a tricky problem and has been discussed controversially in the literature. When considering the problem in the context of the statistical evaluation of spatio-temporal spike patterns using SPADE, we observed that the spike reduction in the UD surrogates in combination with binarization of the spike data leads to fewer pattern occurrences in the surrogate data compared to the original data, which in turn leads to an overestimation of pattern significance in the original data (Fig 7). The ultimate consequence of this problem is the occurrence of false positives. Fortunately, SPADE is a modular method: different types of surrogates can be used, while the mining algorithm and testing steps stay identical. For this reason, we were able to analyze the same data sets by using different surrogates, to then compare the results.

Thus, using SPADE we tested all six surrogate techniques with respect to false positives on non-stationary ground truth data. These were based on two experimental sessions of motor cortex of macaque monkey involved in a reaching-and-grasping task (Riehle et al., 2013; Brochier et al., 2018). We stress the importance of generating test data that are very similar to experimental data, in order to closely model all features that typically lead to complications when trying to robustly estimate the null-hypothesis of conditional independence given the firing rates (Grün, 2009). Such realistic data serve as ground truth to identify the strengths and weaknesses of the tested surrogate techniques. In this case, we modeled both experimental sessions as PPD and Gamma processes, with firing rate modulations, dead-times, and regularities estimated for each neuron from the original data (Fig 8A). The analysis of these data with SPADE led to a large number of FPs when employing UD. However, all other surrogate techniques showed a considerably low number of FPs. A minimal number of false positives is to be expected, as it is inherent to any statistical test. Thus, we conclude that UD is not appropriate for its employment within the context of the SPADE analysis, whereas all other techniques can be considered valid.

Finally, we analyzed experimental data from Brochier et al. (2018). UD in this context leads to a large number of detected patterns (Fig 10). Given the results obtained from the previous sections, analytically and through simulations, we consider these patterns as putative false positives (taking into consideration that in the case of experimental data we have no ground truth at hand). In contrast, employing the other surrogates the number of patterns detected is much smaller than using UD. Still, the number of patterns is larger than for the analysis of the independently generated artificial data with these surrogates. Given also our previous results, we consider the patterns detected in the experimental data by the alternative surrogate techniques as significant, i.e., the patterns do not result from any overestimation of the significance. This is confirmed by the fact that the patterns, retrieved for UDD, JISI-D, ISI-D, WIN-SHUFF, and TR-SHIFT, show almost identical participating neurons, lags, and occurrence numbers. Hence, we conclude that the different surrogates, even though they move the spikes in different ways, lead to an almost identical significance level.

We come to the conclusion that UDD, JISI-D, ISI-D, WIN-SHUFF, and TR-SHIFT are appropriate for the detection of spatio-temporal spike patterns in the SPADE analysis, and UD is not. Furthermore, we suggest TR-SHIFT as the surrogate method of our choice for the SPADE analysis, because it is a technique that 1) is easy to explain and to implement, 2) reflects more closely the hypothesis of temporal coding, 3) reproduces exactly the most relevant statistical features of a spike train (Table 1 and Fig 4), 4) is as conservative as the other methods that we propose, and, 5) employs fewer parameters than the other techniques with the same performance.

Of course, the employment of surrogates is not only restricted to the context of a SPADE analysis but was used in other studies for the evaluation of correlations (Gerstein et al., 1978; Hatsopoulos et al., 2003; Pipa and Grün, 2003; Pipa et al., 2003; Pazienti et al., 2007; Pipa et al., 2007; Maldonado et al., 2008; Smith and Kohn, 2008; Pazienti et al., 2008; Grün, 2009; Harrison and Geman, 2009; Louis et al., 2010a; Dann et al., 2016; Torre et al., 2016). Still, the choice of a particular surrogate technique has to be done appropriately and cautiously case by case. Not only because the statistical test might produce false positives (or false negatives), but also because the concrete null-hypothesis distribution represents the model that is to be falsified. The degree of how conservative or liberal the statistical analysis can be, through the choice of the surrogate technique, becomes then not only a feature of the test but more a scientific question per se regarding neural coding.

Several studies have already investigated the impact of different surrogate techniques in the context of spike time correlations. For example, in Louis et al. (2010b), the authors evaluated the influence of surrogate techniques on cross-correlation analysis of two parallel spike trains; in Grün (2009) and Louis et al. (2010a), the focus was on the effect of surrogate techniques on synchronous events in the context of the Unitary Events analysis (Grün et al., 2002a,b; Pipa and Grün, 2003; Pipa et al., 2003, 2007, 2013). Due to the results of these studies, we have concentrated on surrogate techniques that preserve the firing rate profile of the original neurons. Methods such as spike train randomization (within single trials; Grün et al., 2003), spike exchange (across neurons or trials; Harrison et al., 2007; Smith and Kohn, 2008), ISI shuffling (within and across trials; Ikegaya et al., 2004; Masuda and Aihara, 2003; Nádasdy et al., 1999; Rivlin-Etzion et al., 2006), spike shuffling across neurons (within-trial; Ikegaya et al., 2004; Nádasdy et al., 1999) do not fulfill our requirements (Grün, 2009). Other methods are designed to preserve the auto-correlation of a spike train, with the assumption of stationarity and the Markov property of a process (Ricci et al., 2019; Perinelli et al., 2020). Some studies have already shown evidence of problems arising from the application of uniform dithering, such as the non-preservation of the ISI distribution (Louis et al., 2010a), in particular in the case of the Poisson process (Platkiewicz et al., 2017), but not in the context of multiple parallel spike trains, or in the context of binarization. We extended previous similar comparative studies of surrogate techniques (done only for pairwise correlations; Louis et al., 2010a) to the context of precisely timed higher-order spike correlations. The relevance of this step is sensible, as it has been argued that the processing of information may be reflected in the presence of delayed higher-order correlations in parallel spike trains, in particular, within the context of the synfire chain model (Abeles, 1991; Diesmann et al., 1999; Izhikevich, 2006; Bienenstock, 1995; Oettl et al., 2020). Given our present state of ignorance concerning the detailed computational functioning of the brain, the use of statistical methods like SPADE seems to be almost the only way to find out whether the detailed, precisely timed coordination of several individual neurons is important for information processing beyond the more global temporal covariations of neural activity that move between cortical layers and across cortical areas.

We have started to apply SPADE to a large set of experimental data to investigate the presence of spatio-temporal patterns in and their relation to behavior, and first encouraging results are shown in Fig 10.

## Author Contributions

Conceptualization, A.S., P.B., G.P., S.G.; Data generation, A.S., P.B.; Data analysis, A.S., P.B.; Software implementation, A.S., P.B.; Visualization, A.S., P.B.; Writing, A.S., P.B., G.P., S.G.; Project administration, S.G.; Funding acquisition, S.G. All authors have read and agreed to the published version of the manuscript.

## Acknowledgments

This project was funded by the Deutsche Forschungsgemeinschaft (DFG, German Research Foundation) - 368482240/GRK2416; by the European Union’s Horizon 2020 Framework Programme for Research and Innovation under Specific Grant Agreement No. 785907 (Human Brain Project SGA2), No. 945539 (Human Brain Project SGA3); by the Helmholtz Association Initiative and Networking Fund under project number ZT-I-0003; and by the Joint-Lab “Supercomputing and Modeling for the Human Brain”. We thank Sebastian Lehmann (SBC) for the help in the design and development of the graphics.

